# Quantitative dissection of color patterning in the foliar ornamental Coleus reveals underlying features driving aesthetic value

**DOI:** 10.1101/2021.01.11.426252

**Authors:** Mao Li, Viktoriya Coneva, David Clark, Dan Chitwood, Margaret Frank

## Abstract

- Coleus is a popular ornamental plant that exhibits a diverse array of foliar color patterns. New cultivars are currently hand selected by both amateur and experienced plant breeders. In this study, we reimagine coleus breeding using a quantitative color analysis framework.
- Despite impressive advances in high-throughput data collection and processing, complex color patterns remain challenging to extract from image datasets. Using a new phenotyping approach called “ColourQuant,” we extract and analyze pigmentation patterns from one of the largest coleus breeding populations in the world.
- Working with this massive dataset, we are able to analyze quantitative relationships between maternal plants and their progeny, identify features that underlie breeder-selections, and collect and compare consumer input on trait preferences.
- This study is one of the most comprehensive explorations into complex color patterning in plant biology and provides new insights and tools for exploring the color pallet of the plant kingdom.

## Introduction

Coleus (*Coleus scutellarioides*) is a common ornamental bedding plant that is bred for its brilliant and diverse foliar color patterning (Bailey, 1924; Pedley & Pedley, 1974; Paton *et al.*, 2018, 2019). Wild relatives in the Coleus genus harbor a small degree of variegated pigmentation that has been expanded into distinctive new cultivars that harbor complex variegation patterns through successive rounds of hybridization and selection (Suddee *et al.*, 2004). The prevalence of Coleus in gardens and urban landscapes around the world is a testament to the unique aesthetic capacity of this species (Rogers, 2008). With over 500 cultivars on the market, and new ones added each year, coleus represents one of the largest and most diverse examples of pigmentation patterning within a single species.

Advances in plant phenotyping have revolutionized how humans interact with botanical traits (Fahlgren *et al.*, 2015; Gehan & Kellogg, 2017; Gehan *et al.*, 2017; Li *et al.*, 2018b; Prunet & Duncan, 2020; Amézquita *et al.*, 2020). High throughput data collection has enabled rapid agricultural trait selection (Singh *et al.*, 2019; Shakoor *et al.*, 2019; Ibba *et al.*, 2020), early detection and management of disease (Mutka & Bart, 2014; Shakoor *et al.*, 2017), and large-scale 2-dimensional morphological analyses (Li *et al.*, 2018a). Penetrating high-resolution imaging technologies, such as X-ray CT and laser ablation tomography have also made complex, three-dimensional topologies accessible (Chitwood *et al.*, 2019; Li *et al.*, 2019b, 2020a; Prunet & Duncan, 2020; Amézquita *et al.*, 2020; Vanhees *et al.*, 2020). Despite these enormous advances, rapid phenotyping for complex color patterning remains a major hurdle in High Throughput (HTP) analysis. Indeed, the majority of color phenotypes expressed in plants are typically uniformly expressed (for example, monochromatic leaves (Gehan *et al.*, 2017) and berries (Underhill *et al.*, 2020)), un-patterned in their expression (for example, lesions (Arnal Barbedo, 2013; Gobalakrishnan *et al.*, 2020; Xie *et al.*, 2020)), or have highly predictable patterns (for example, nectar guides). These color phenotypes are readily extractable using existing image processing approaches that are not suited for the complex suite of color patterns represented in our coleus population (Arnal Barbedo, 2013; Gobalakrishnan *et al.*, 2020; Xie *et al.*, 2020)). Here, we address the need for enhanced tools to extract and analyze complex patterns. In this study, we map out pigmentation values as three-dimensional point clouds in Lab color space, extract the continuous distribution of color using Gaussian density estimation (Li *et al.*, 2019a), dissect color patterns based on pigmentation position on two dimensional leaves, quantify bilateral symmetry for shape and color, and separate shape from color using thin plate spline deformation.

Given the prominence of Coleus in the gardening marketplace, and the vast diversity of pigmentation patterns that are exhibited within Coleus breeding populations, Coleus as a breeding system serves as an ideal platform for testing this new, quantitative approach for HTP color phenotyping. In this study, we develop a pipeline to extract quantitative descriptors for foliar pigmentation patterns from one of the largest Coleus breeding populations in the world (n > 32,800 plants). We are able to extract the distribution of all existing pigmentation patterns presented within this massive breeding population, quantify maternal plant-progeny pigmentation relationships, and identify aesthetic features that are associated with increased value from the perspective of the breeder as well as the general public. This work is built on a powerful study system, and provides a new framework for approaching complex color phenotyping. This work has direct implications for investigating color features in both ornamental plant breeding and ecological systems, where pigmentation patterns play an important role in influencing how plants interact with humans, pollinators, and herbivores.

## Methods

### Coleus population, sampling, and image processing

We collected and sowed 50,000 Coleus seeds from 133 open-pollinated mother plants in early January, 2015 in Gainesville, FL. We organized the seedlings into families based on their maternal parents, grew the plants for five weeks and then selected ~2,000 individuals as potential new cultivars based on their foliar color patterning and branching architecture in mid-February. Next, we harvested the youngest fully expanded leaf from each plant between 5-6 weeks of age, and imaged the leaves on Epson Perfection V550 Scanners with Kodak KOCSGS color separation guides included for color calibration (Supplemental Fig 1; data available here: Zenodo.org 10.5281/zenodo.4421754). We performed color analysis using our open-access software program called ColourQuant (Li *et al.*, 2019a); software available on github: github.com/maoli0923/ColourQuant). Briefly, we adjusted the RGB color balance on each scan by a white balance method so that the white swatch in the Kodak KOCSGS color separation guide is pure white, to ensure that scanners were not biasing the color data. Next, we segmented the leaves from the background by converting the RGB matrix into hue-saturation-value (HSV) format. Since most background pixels are grey in HSV, this was used to set a threshold (e.g. S>0.15) that separates grey values from true leaf values. We then used the binary leaf silhouettes to extract the leaf color data by setting the background to pure white. We manually adjusted the thresholding for leaves that could not be automatically extracted due to shadows in the scan, and removed outliers from the sample set, including leaves that were overlapping on the scanner, very small, or broken.

### Color pattern analysis

To extract quantitative color distribution information, we converted the leaf color matrices from RGB to CIELAB (L*a*b*) color, which is a continuous color space that consists of three descriptors: L* = “lightness,” a* = “green to magenta,” and b* = “blue to yellow.” We studied the distribution of mean and variance for L*, a*, b* color values across the leaves by first calculating the average value and variance of L*, a*, and b* for each leaf (i.e. “mean L”, “mean a”, “mean b”, “variance of L”, “variance of a”, and “variance of b”), and then plotting histograms and boxplots to show the overall mean and variance distributions for all leaves in the breeding population. Next, we treated the 3D Lab color matrices as 3D point clouds, which enabled us to extract color distribution and frequency information for each leaf.

The mean and variance of Lab values roughly describes the color for each leaf. However, in order to compare the distribution and frequency of Lab values across the leaves, we applied a Gaussian density estimator (GDE) to the Lab point cloud. GDE is a function defined on 3D space, providing a robust and direct density estimate from the point cloud data. To reduce computational complexity, we restricted the domain of the GDE function to a fixed bounded cuboid. The GDE descriptor alone captures statistical color frequency, not spatial patterning. To capture spatial color information, we segmented the leaves into distinct zones based on normalized pixel distances: “border” – defined as the outer 15% of pixels from the leaf boundary to the centroid, “center” – defined as the inner 75% of pixels from the centroid to the boundary, and “full” – defined as the entire color matrix. The distance between any two leaflets is calculated with the following equation:

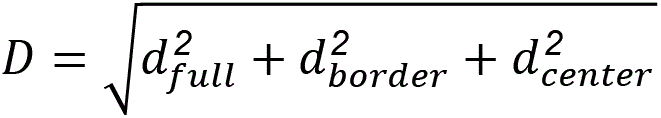

where *d* represents the L_2_ distance between GDE functions for each corresponding zone. With this calculation, the pattern difference between two leaves is determined by their degree of similarity across all three zones. For pairwise distances, we used multidimensional scaling (MDS, similar to a PCA) to project the data in a lower dimensional space, which allows us to capture the major features that contribute to pattern variation. These methods and the supporting software for this approach can also be found in the publication by (Li *et al.*, 2019a) (https://github.com/maoli0923/ColourQuant).

To quantify the degree of mirror symmetry for each leaf, we first marked a bilaterally symmetric line by placing two landmarks, one at the proximal point (petiole) and another at the distal point (leaf tip). These landmarks were then used to partition the leaf into longitudinal halves that could be directly compared to one another. We used two methods for quantifying mirror symmetry. First, we performed a general measure by comparing the differences in left and right color distributions (using GDE functions), and second, we measured the degree of bilateral shape symmetry by overlaying the left and right halves of the leaf and computing the percentage of pixels that fail to overlap. Notably, smaller values reflect a lower degree of asymmetry.

### Quantitative analysis of maternal-offspring pigmentation relationships

To calculate the phenotypic distance between maternal plants and their progeny, we divided the distance between each maternal leaf and the leaves of its progeny by the distance between the maternal leaf and all of the leaves in the breeding population. To investigate how maternal color and color complexity influence these color traits in the progeny population, we calculated the mean and variance of L*, a*, and b* for the maternal leaves and their offspring and then computed the variance of those traits across the offspring within each family (e.g. named as “variance of family mean L”, and “variance of family L variance”).

### Quantifying aesthetic features of selected plants

We calculated the influence of breeder selection on color and shape symmetry, as well as pigmentation L*, a*, and b* values, by comparing the probability distribution for each value in the entire breeding population with the probability distribution in the selected population. Two sample T-tests accounting for uneven sample sizes were used to calculate the significance of selection on each color parameter.

### Public preferences for coleus colors independent of shape

To investigate color preferences amongst the general public, we created a survey based on the major sources of variation for leaf color patterning. First, we separated shape from color patterning by deforming the leaves into uniform circles using thin-plate-spline (TPS) interpolation, followed by centering and normalizing the circles into the same position and size. Next, we rotated each circularized leaf so that the first landmark (near the base) is on the negative half of the y axis (x=0) and the tip is on the positive half. We resized each circular leaf image to be 70×70 dimension and reshaped the pixel L*a*b* colors into a long (12150 dimension) vector. To calculate the main sources of variance, we performed a principal component analysis on the long vectors of the circularized leaves and created a survey using google forms where public volunteers were asked to select their preference of eigencolors for the top 8 principal components (PCs). For kth PC, the eigencolors are represented by ± x standard deviation along PC axis, where x=3+(k-1)*0.5, this produced more distinct color variants for the survey participants. We distributed the survey using a dedicated Twitter account (@ColeusColours), and then plotted the responses from all of the survey participants (n=172) and reconstructed the composite preferred leaf based on the responses.

## Results

### New coleus breeding population

Coleus is one of the most diverse species with regards to leaf pigmentation patterning in the world. Brilliant new coleus cultivars harboring novel leaf color and shape phenotypes can be generated using a recurrent mass selection approach. In this study, we took advantage of a very large coleus breeding population in order to explore the full spectrum of possible pigmentation patterns and their influence on breeding processes. We used 133 open-pollinated elite coleus lines that exhibit a wide range of existing color and shape phenotypes (Fig 1A-B) to generate a large population that harbors novel pigmentation combinations. To capture these new combinations, we planted over 32,000 F1 progeny, and imaged their leaves on high-resolution color scanners (supplemental Fig S1). Color data are typically recorded as a composite of discrete Red, Green, and Blue (RGB) values that range from 0-255. We transformed our RGB data into the continuous Lab color space, which we then treated as a three-dimensional point cloud and extracted quantitative pigmentation data using a Gaussian density estimator (GDE) function (Fig 1C). A GDE function is a smoothed version of a histogram; it estimates data density by summing all of the normal distributions, which are placed on each data point. Higher values are produced from regions with more data points, while lower values are produced from regions with sparse and/or noisy data, thus making the function robust.

**Figure 1:**
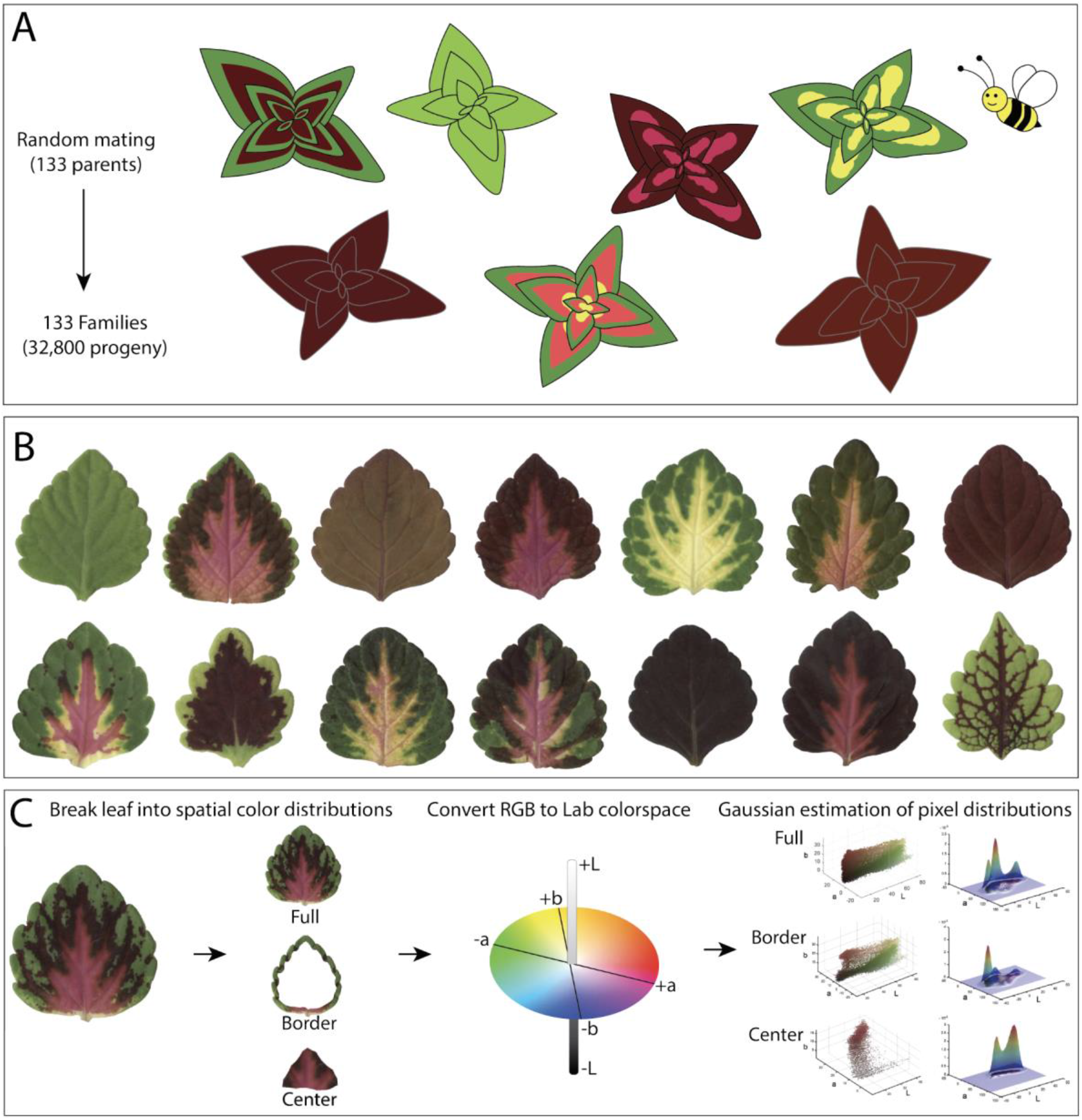
Experimental design, high throughput sampling, and color analysis. (A) 133 field-grown parents were randomly mated by pollinators, seeds were collected from each maternal plant, sown in progeny family blocks and grown for 5-6 weeks in a greenhouse; (B) One fully-expanded leaf was harvested and scanned from each plant in the population; (C) Color thresholding was used to isolate binary masks for each leaf. Discrete RGB color matrices were converted to the continuous Lab color space, and color matrices for each leaf were spatially separated into segments: “full” – defined as the entire color matrix, “border” – defined as the outer 15% of pixels from the leaf boundary to the centroid, and “center” – defined as the inner 75% of pixels from the centroid to the boundary. A Guassian density estimator was used to extract quantitative pigmentation data (only 2D Gaussian density estimator was shown for visualization).

To visualize the CIELAB (L*a*b*) color space within our breeding population, we plotted the mean values of L* (lightness), a* (green-to-magenta), and b* (blue-to-yellow) that were extracted from each leaf. The majority of leaves within the population skewed towards darker (lower) mean L values (Fig 2A). Mean values for a* spanned from magenta-to-green, but were more heavily concentrated towards the magenta/maroon half of the range (Fig 2B), and mean values for b* were almost exclusively in the positive range, and were strongly concentrated towards yellow rather than blue values (Fig 2C). While this approach provides an estimate of mean color distributions, it fails to capture color patterning within the population (Fig S2). There are three discrete regions that can be used to generally describe that vast majority of variegation patterns in coleus: the area surrounding the veins, the leaf border, and the leaf center. We applied a Gaussian density estimator function to 3-dimensional point clouds of the border (15% of pixels from the leaf boundary) and center (75% of the pixels from the centroid) regions of the leaf (Fig 1C). Venation varies considerably from leaf-to-leaf, and thus it is challenging to consistently extract this value from a large population, so we did not consider the contribution of variegated venation for this study. Our isolated border and center regions differed significantly from the full variance of L*a*b* values (P ranged from 7.75 e-04 to < 2.23 e-308), indicating that these regions exhibit distinct color patterns (Fig 2D). Multidimensional scaling (MDS) can be used to extract the main sources of variance within complex datasets. To investigate the variance in color patterning within our population, we generated MDS plots from the GDE function distance for the full leaf (Supplemental Fig S3A), border (Supplemental Fig S3B), center (Supplemental Fig S3C), and composite full leaf plus border plus center distance (Fig 2E-H). We have superimposed example leaves on top of the plot to illustrate the major color differences that are represented within the population (Fig 2F, H, and S3).

**Figure 2:**
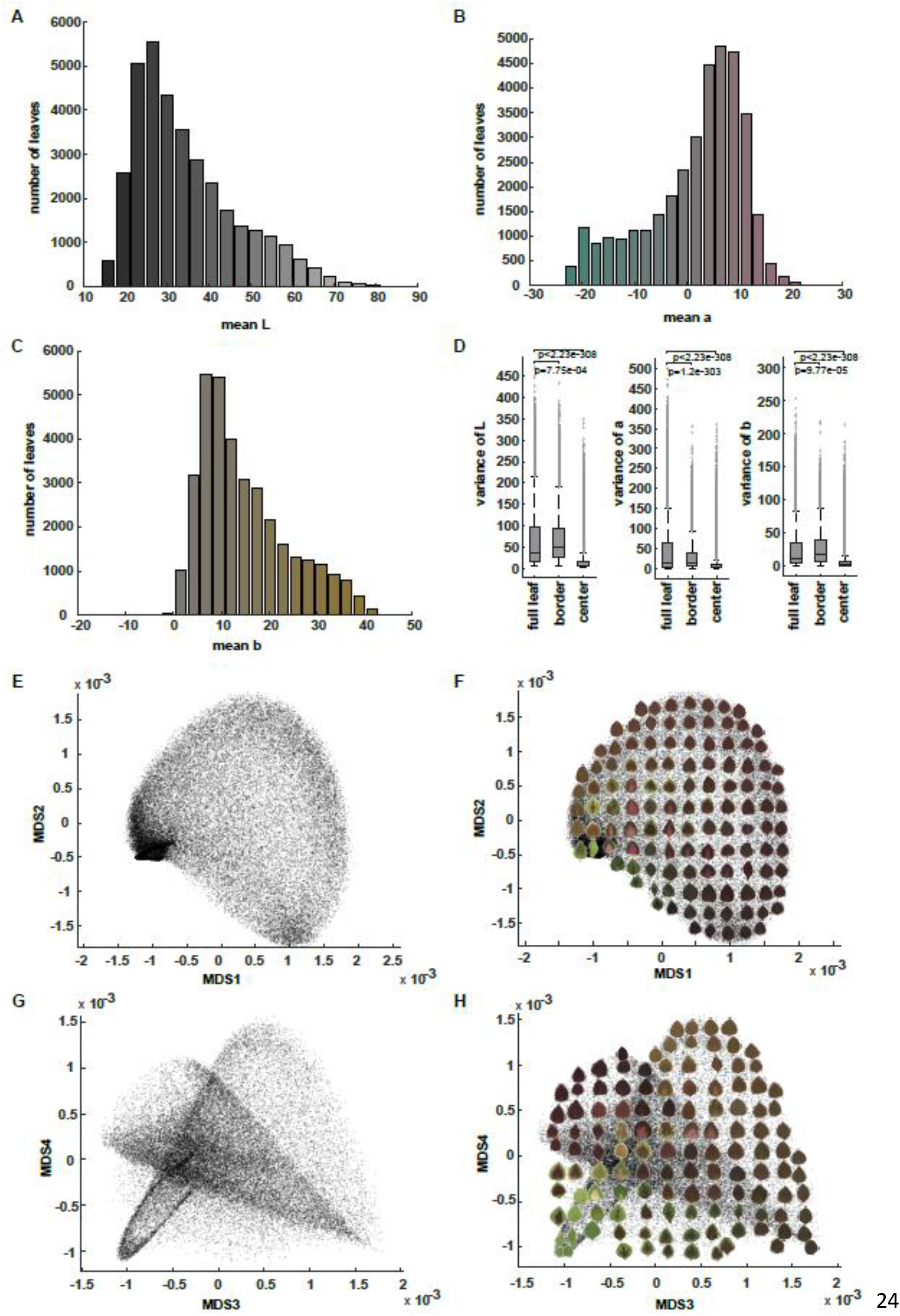
CIELAB (L*a*b*) color distribution. **(A)** The histogram of mean L (lightness) values of the studied coleus population. The color for each bar corresponds to the Lab color with L value at x axis, a=0 and b=0; **(B)** Histogram of mean a (green to magenta) values. The color for each bar corresponds to the Lab color with a value at x axis, L=50 and b=0; **(C)** Histogram of mean b (blue to yellow) values. The color for each bar corresponds to the Lab color with b value at x axis, L=50 and a=0; **(D)** Boxplot of the variance of L, a, and b for full leaf, border, and center. The “+” signs mark outliers that are more than 1.5 interquartile ranges above the upper quartile or below the lower quartile for each box, the central line indicates the median, top and bottom edges of the box indicate 25th and 75th percentiles. Whiskers extend to the most extreme non-outliers of the data. P-values for full leaf versus border, full leaf versus center are also shown using paired sample t-test; **(E)** and **(G)** Multidimensional scaling (MDS) plot (MDS1 vs MDS2 in (E) and MDS3 vs MDS4 in (G)) for the pattern difference defined by the difference of Gaussian density estimator in 3D Lab colorspace across full leaf, border and center; **(F)** and **(H)** The same MDS plots shown in (E) and (G) but with example leaves superimposed to provide visual examples of the data distribution.

The sub-sample border and center plots provided poor separation of the major pattern classes within the population (Supplemental Fig S3B-C). For example, green bordered leaves with maroon centers are distributed in multiple locations across the border and center MDS plots (Supplemental Fig S3B-C). The full leaf plot performed much better with regard to pattern separation compared to the sub-sample plots; however, it still failed to produce distinct groupings for detailed pattern differences. For instance, pink and maroon center variegation patterns are intermixed with solid maroon leaves in all 4 dimensions of the full leaf MDS (Fig S3A). The composite plot, on the other hand, accounts for both global and isolated center and border pigmentation values, and thus was able to resolve distinct pattern groupings (Fig 2E-H). The first dimension clearly separates the population along the green-to-magenta divide (the a* value of the L*a*b* color space), while the second dimension separates the population from darker (towards the bottom) to lighter L* pixel distributions (Fig 2F). In the third and fourth dimensions, five major patterns are resolved: solid orange in the upper right, solid deep purple in the upper left, solid green in the lower left, solid maroon in the lower right, and several sub-populations of variegated patterns in the lower left and center (Fig 2H). In the lower left corner of MD 3 and MD 4, we were able to resolve most of the variegated patterns into subpopulations based on center and border features, for example wide maroon centers with thin green borders, light pink centers with green, maroon, or orange borders, yellow/white centers with green borders, and even deep purple venation on green leaves. There are, however, two pigmentation patterns that we failed to isolate in our composite plot. First, are leaves that have both relatively small central pigmentation regions and low contrast between the border and center colors, and second, are leaves with random green and purple sectors whose patterns were most likely generated by active transposons (Tilney-Bassett & Others, 1986; Frank & Chitwood, 2016). Overall, this composite MDS approach performed very well with regard to separating the population into major pattern groups.

### Maternal phenotypes influence the phenotypic distance of their progeny

The vast majority of brilliant new coleus color patterns result from the spatially regulated production of anthocyanin (purple and pink pigments) and loss of chlorophyll (white and yellow pigments). Classic genetic analyses indicate that purple pigmentation is controlled by a single dominant allele, while loss of chlorophyll pigmentation resulting in yellow/albino phenotypes results from a recessive allele (Boye & Rife, 1938; Rife, 1948). These studies were carried out in simplified phenotypic and genetic backgrounds; however, they provide a basic framework for interpreting the color relationships within our large breeding population. To address patterning relationships within our population of 32,000+ individuals, we quantified the relative distance between maternal plants and their progeny and visualized it in multidimensional scaling (MDS) space (Fig 3). It is important for us to note that our population was generated using an uncontrolled, open-pollination design; honeybee hives were brought into the field to ensure pollination and promote outcrossing amongst the maternal plants. In our field setting, it is impossible to track the male half of the parental equation without the developing genotype-specific molecular markers, so we are only analyzing maternal-to-progeny relationships. Another limitation that we cannot exclude from this experimental design is the potential bias that leaf patterning can have on pollinator behavior, and while we cannot assume true random mating within this context, we have reason to believe that pollination behavior is close to random based on the fact that coleus flowers tend to be highly conserved with respect to their morphology and color. Thus, they are likely equally attractive to our honey bee pollinators.

**Figure 3:**
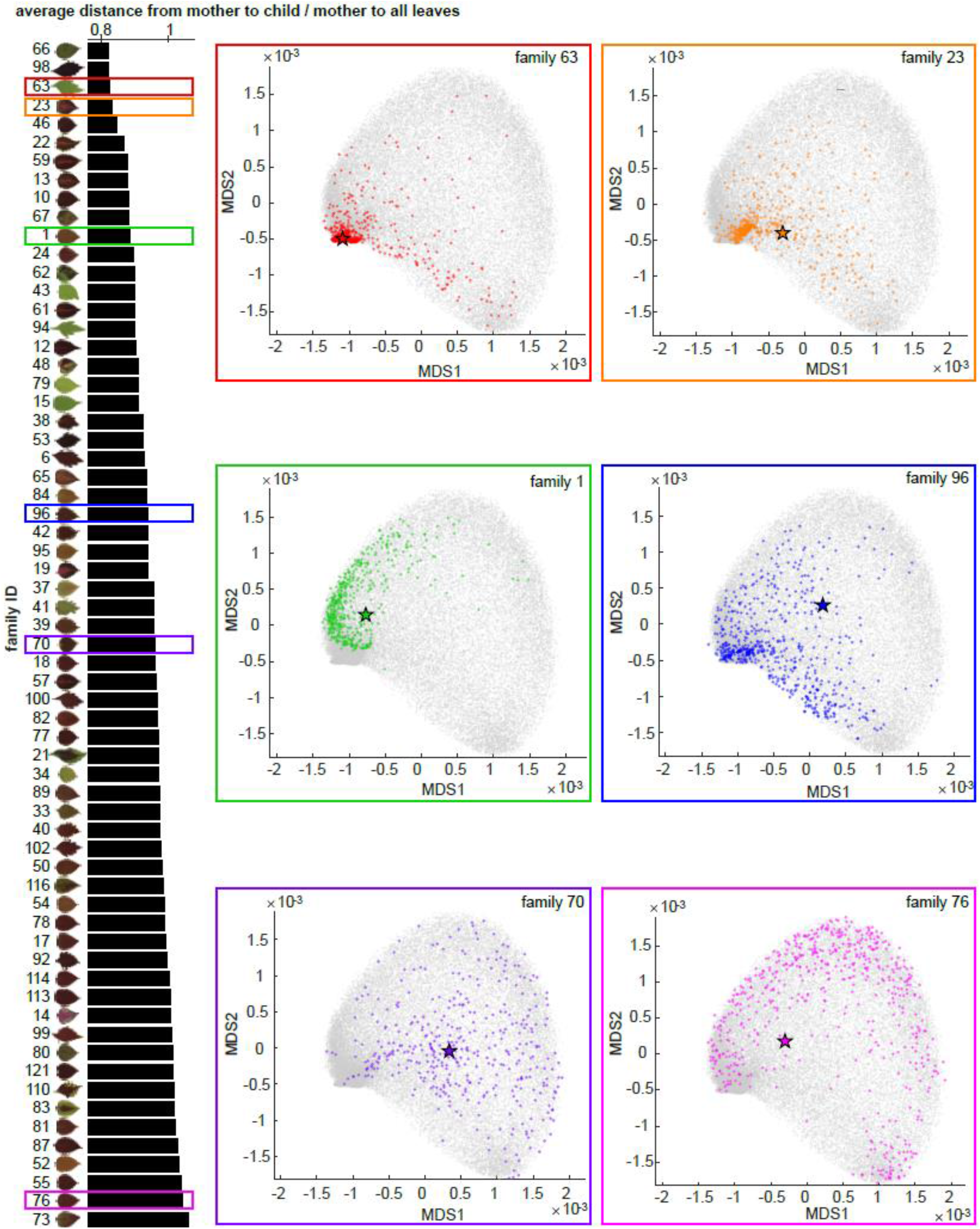
Maternal Plant-Progeny relationships. On the left panel, each bar shows the average distance from maternal plants to progeny divided by the average distance from maternal plants to all leaves (x-axis) for each progeny family (y-axis) superimposed upon the scan of the maternal plant. On the right panels, there are six MDS plots (MDS1 vs MDS2) from six progeny families as examples with different colors correspond to the families highlighted in the same colored rectangles on the left panel. On each MDS plot, grey dots show all leaves, colored stars represent the maternal plants, and colored dots are the progeny.

We identified a few clear trends from our mother-child analysis. First, brighter maternal plants (high L*) tend to produce progeny with a greater variance of pixel brightness (Fig. S4A). This is exemplified by the progeny in families 79 and 43. We also observed that green maternal plants (low a*) tend to produce progeny that exhibit a large variance between green and magenta (Fig. S4B, for example, the progeny in families 43 and 94). This is logical, given that purple and magenta pigments have been linked to dominant alleles, and thus would be expressed in F1 crosses with purple/magenta pollen donors. Along similar lines, yellow maternal plants (high b*) tend to produce progeny that express a large variance in the yellow-to-blue color range (Fig. S4C, for example the progeny in families 79 and 29). Again, this follows the logic that yellow pigmentation is a recessive trait, and thus color patterning in the F1 generation is more likely to exhibit paternal phenotypes. We found that maternal plants with complex color patterning (high variance of L*, a*, or b*) tend to produce progeny with larger variance in their complexity (Fig. S4D-F, for example the progeny in families 22 and 23), which results in more diverse color patterns. Surprisingly, we only saw a minor trend for green versus purple maternal plants being closer versus farther away (respectively) from their progeny in phenotypic space. The majority of green leafed maternal plants fall on the top half of the phenotypic distance plot (e.g. smaller distance, for example, the progeny in families 63, 43, 94, 79, and 15), while purple maternal plants are distributed across the phenotypic spectrum (Fig 3).

### Bilateral symmetry for color and shape are strongly correlated with the selection of new cultivars

New coleus cultivars are hand selected based on the visual identification of target traits, through a process that is frequently referred to as selection via “the breeder’s eye” (Fasoula *et al.*, 2019). Our experienced coleus breeder identified approximately 2,000 selected lines from the population to carry forward for potential cultivar development. A long-standing theory posits that symmetry is positively correlated with aesthetic value (Birkhoff, 1933). To investigate the influence of color and shape symmetry on our breeding process, we tested whether our selected population deviated significantly from the total population with regard to color and shape symmetry, as well as mean Lab distributions (Fig 4).

**Figure 4:**
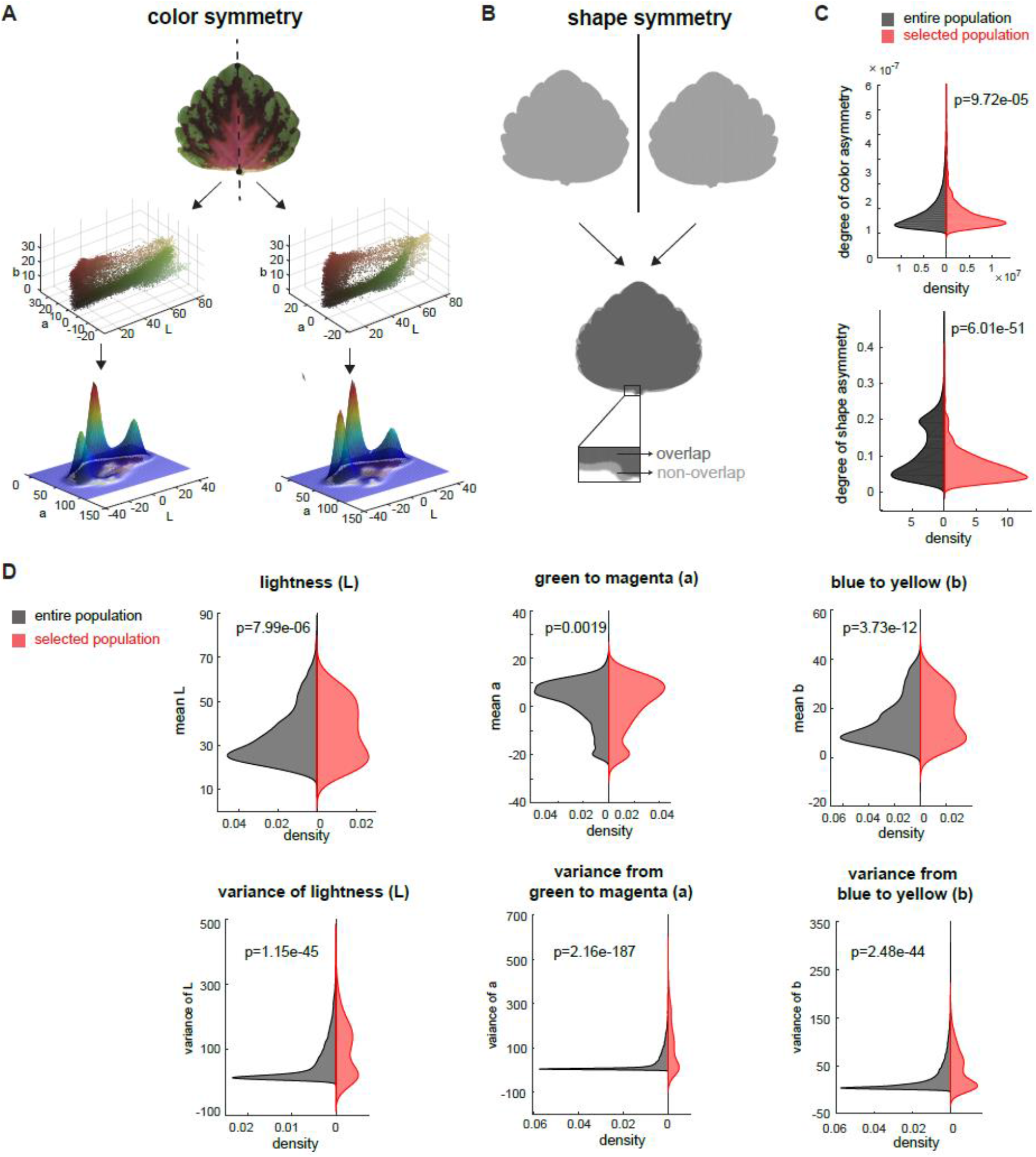
Influence of color and shape on cultivar selection. **(A)** Mirror symmetry of color: Partitioning of each leaf into left and right halves (top panel), convert each part into 3D point cloud in Lab color space (middle panel), and calculate the 3D Gaussian density estimator (lower panel, only shows 2D Gaussian density estimator for visualization); **(B)** Mirror symmetry of shape: flip the leaf horizontally (top panel), measure the non-overlapped area (lower panel) and calculate the percentage of non-overlapped area over the leaf area; **(C)** Distribution of degree of color asymmetry (top panel) and shape asymmetry (bottom panel) for entire population (in black) and selected population (in red); **(D)** Distribution of mean L (top left), mean a (top middle), mean b (top right), variance of L (bottom left), variance of a (bottom middle), and variance of b (bottom right) for entire population (in black) and selected population (in red). Significance was measured using a two sample t-test for uneven sample sizes.

To quantify the degree of mirror symmetry within leaves from the selected versus total population we manually partitioned every leaf into left and right halves by drawing a line from the tip to the base of the leaf. We then quantified color asymmetry by comparing the Lab Gaussian distributions between the left and right halves (Fig 4A), and shape symmetry by folding binary leaf silhouettes along the midline and calculating the percentage of non-overlapped pixels (Fig 4B). Our two sample T-tests between the selected and total population showed very strong statistical support for both color and shape symmetry playing a significant role in influencing the selection process (p-value 9.72e-05 for increased color symmetry, and p-value=6.01e-51 for increased shape symmetry in the selected population; Fig 4C).

To determine if specific color features correlated with cultivar selection, we tested whether the selected pool differed from the total population with regard to independent components of the Lab color space (Fig 4D). Interestingly, the selected pool deviated significantly from the full population with regard to both the mean and variance for each of the three Lab color components (Fig 4D). Comparative plots of mean Lab space for the total population (in gray) and selected pool (in red) clearly show that the source of divergence between these two populations comes from an accentuated bimodal distribution on either end of the spectra within the selected pool, indicating that the breeder is selecting along the extremes of the color space. For example, within the L spectrum (the light-to-dark spectrum), the enriched bimodal distribution, reflects strong selection for both bright and dark (deep colored) pixel values (p-value for mean L = 7.99e-06; Fig 4D). Furthermore, our analysis revealed significant divergence in the distribution of selected versus total population values for variance within the Lab space (p-values for L=1.15e-45, a=2.16e-187, b=2.48-44). Again, graphs for the selected pool have strong bimodal distributions for all three Lab spectra indicating that there was selection for varieties with either high color contrast or uniform (solid color) patterning (Fig 4D). In contrast, the total population graphs are concentrated around a single mean peak (Fig 4D). Taken together, this analysis demonstrates how the “breeder’s eye” reshaped the selected pool to significantly enhance mirror symmetry for both color and shape, and concentrate the cultivars with either high color contrast or complete color uniformity. Notably, this analysis accounts for the first round of selection where a high level of variability concentrated around both commercial targets and novel aesthetic traits are maintained. Approximately 6-8 of the plants from this large selection pool are taken through the commercialization process.

### Public survey shows strong overlap between public preferences and breeder selection

Once we established the quantitative color structure for our breeding population, we explored how the existing coleus color space matched with public color preferences. To do this, we created a pilot survey that was openly distributed using a dedicated Twitter account (@ColeusColours). To avoid the confounding influence of leaf shape on color preference, we standardized the leaf orientation based on the bilateral symmetrical line and deformed our leaf shapes into circles using a thin plate spline interpolation (Fig 5A), this method smoothly transforms the border shape into a uniform edge with minor distortion of the internal color patterning. Next, we performed a principal component analysis with our circularized leaves (Fig 5B-C) and used the top principal components to construct our survey for color preference. Our survey presented 8 questions that asked the participants to select their preference from the mean and plus or minus a few standard deviations along PC axis (“eigencolors”) for each of the top 8 principal components (Fig 5D). We gathered data from 172 participants, plotted each of their preferences (Fig 5E), and then reconstructed the ideal leaf based on public preferences for the first eight eigencolors with weighted contributions based on the percent variance contained within each PC (Fig 5F). Our results show that participants have a strong preference for very green (responses to PC1 in Fig 5E), very magenta (responses to PC2 in Fig 5E), and leaves with either high contrast color patterns (responses to the contrasting standard deviation extremes in PC3-PC8). The resulting ideal leaf that was reconstructed from the survey data has a high contrast bright green border with internal magenta pigmentation and yellow base (Fig 5F). This ideal leaf not only matches an existing variegated pattern that was resolved in the lower left hand quadrant of MD 3 and MD 4 in our original population analysis (Fig 2H), it is also consistent with the direction of breeding in our selected pool (Fig 4D). This result indicates that even with this small pilot survey, there is strong overlap between public preferences and new cultivar development.

**Figure 5:**
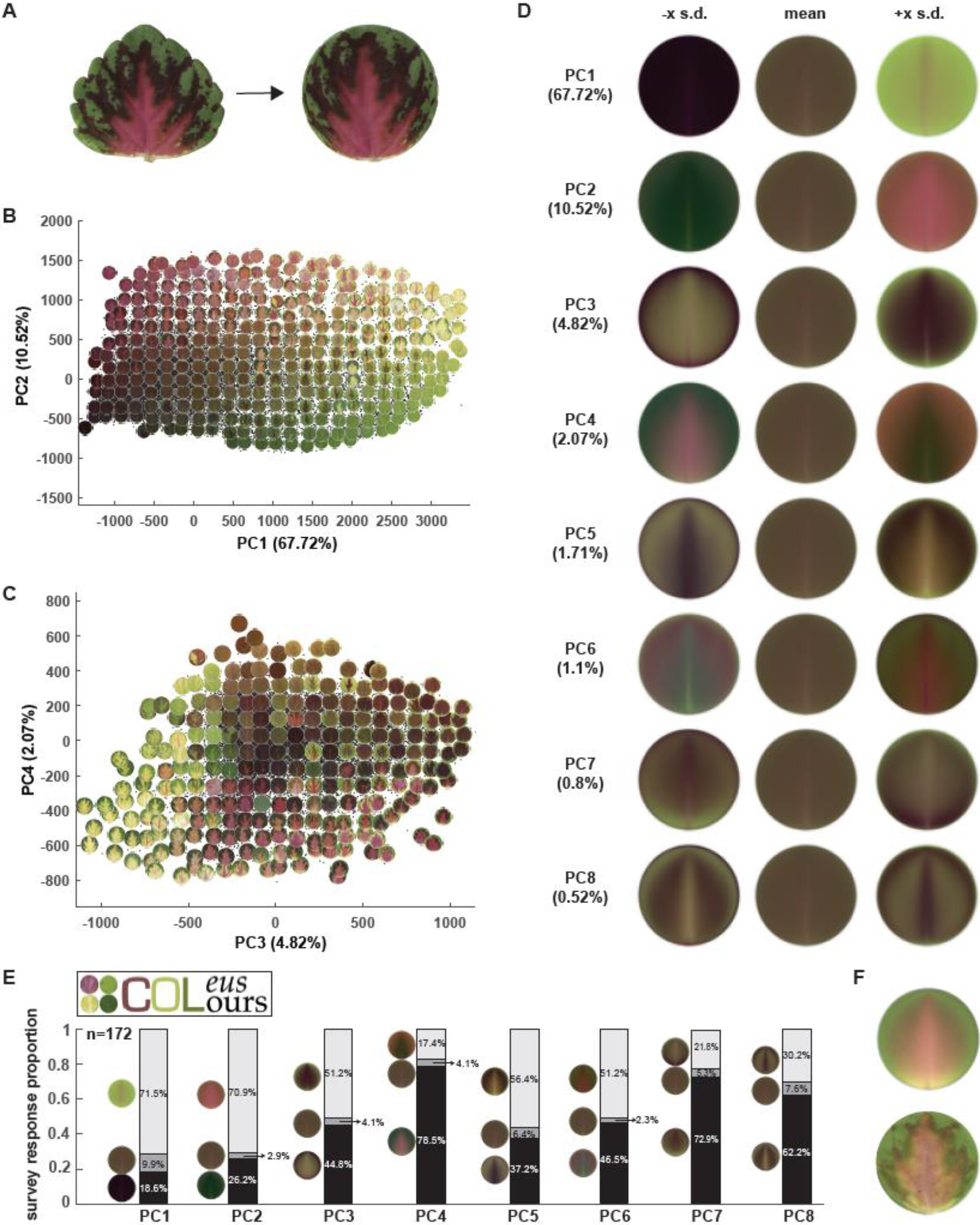
Public survey for color preferences using shape-transformed leaves. **(A)** Deform each leaflet into a disk by thin plate spline interpolation – non-linear deformation into a unit circle; **(B)** and **(C)** Principal component analysis plot superimposed upon some example of leaves (PC1 vs PC2 in (B) and PC3 vs PC4 in (C)) for the pixel Lab values of deformed leaflet; **(D)** Eigencolors for the first eight PCs and the percentage of variance they explained. For PC k, the eigencolor at −x SD and +x SD along PC axis are shown, where x=3+(k-1)*0.5 for better visualization; **(E)** Survey logo (top left) and the survey result from 172 responses; **(F)** Reconstructed pattern (top) and closest real leaf (bottom) from first eight eigencolors with weights guided by the survey response proportion.

## Discussion

High-throughput phenotyping (HTP) has transformed our ability to select and optimize plant traits (Das *et al.*, 2015; Shakoor *et al.*, 2017; York, 2019; Liu *et al.*, 2020). Relative to morphological and architectural phenotypes, approaches for collecting and analyzing color patterns in plants remain limited. Indeed, existing methods of HTP data analysis are not well-suited for the large suite of patterning phenotypes exhibited in ornamental plants, like Coleus. In this paper, we utilize a new approach to address the problem of complex color patterning in a large Coleus breeding population. We partitioned the 2-dimensional leaf into different zones based on morphology and transformed the color data into a continuous, three-dimensional color space, and applied a Gaussian density estimator to extract pixel patterning across space. Using this approach we were able to successfully resolve the major pigmentation patterns contained within one of the largest and most diverse color patterned breeding populations in the world. Historically, these patterns were discussed using qualitative descriptors. By extracting the quantitative features underlying this pattern space, we were able to mathematically analyze relationships between maternal plants and their progeny, identify how aesthetic preferences reshape the color properties of the breeding population, and independently address whether public preferences align with commercial breeding goals.

Our maternal-offspring color analysis may be one of the first times that the inheritance of pigmentation traits has been analyzed through this quantitative lens. We identified quantitative connections between color variance in maternal plants and their offspring that have direct applications for ornamental breeding. For example, breeders looking to increase the range of brightness within their population can start with a brighter parental population; we show that brighter mothers produce offspring that express a wider variance of brightness. Those aiming to increase overall color variation would want to start with parental plants that exhibit complex color patterning, as these mothers produced offspring with the largest variance in terms of pixel complexity. In line with classic genetic studies for coleus that identified purple as a dominant trait and green and yellow as recessive, we found that mothers with pixel concentrations on the green and yellow ends of the spectrum produced offspring that had wider color variation. In essence, recessive color palettes could be considered blank canvases for breeding new pattern variants.

Our analysis of features associated with breeder selection supports long-held theories about aesthetic preferences in humans; aesthetic preference for bilateral symmetry (Birkhoff, 1933) is reflected in the breeding process, where we identified significant enrichment for bilateral color and shape symmetry. Moreover, we found that public preferences for leaves with high color contrast largely agrees with the independent selection process for breeding new cultivars. As mentioned previously, new coleus cultivars are currently sight-selected through a process that involves extensive screening by professional and amateur breeders. The strong quantitative agreement between well-established aesthetic preferences and the breeding process, opens the possibility for automating this first step of cultivar selection. It is not hard to imagine taking this a step further, transforming the cultivar selection process into a customized system. Simple surveys, like the Coleus Colours pilot survey conducted for this study, could help people identify their ideal patterns and automated population screening would match a novel cultivar from the breeding population with the customer. This reimagined breeding approach offers people the personalized experience of designing and naming their own, unique coleus cultivar.

Pigmentation patterns have fascinated scientists for centuries. These visual cues direct plant-pollinator interactions (Leonard & Papaj, 2011; Whitney *et al.*, 2013), fend off herbivores (Lev-Yadun, 2017), and as shown in this study, influence aesthetic value in ornamentals. A simple, yet elegant model involving a reaction-diffusion based mechanism, was famously put forth by Alan Turing to explain the diversity of pattern formation in nature (Turing, 1953). Recent work in the genus *Mimulus* uncovered genetic regulators that fit this Turing-based model, and direct the patterning of nectar guides through a reaction-diffusion interaction between an activator (NEGAN) and its inhibitor (RTO) (Ding *et al.*, 2020). Beyond this specific result, significant progress towards mapping the underlying genetic mechanisms that regulate pigment deposition has been made using diverse floral models. In these systems, an R2R3 Myb, bHLH, and WDR “MBW” transcriptional regulon has been identified as a central regulator for color patterning, controlling both orange carotenoid and purple/red anthocyanin deposition (Sagawa et al. 2016; Ludwig et al. 1989; Albert et al. 2014). In contrast to floral systems, relatively little is known about the genetics of color patterning in vegetative organs; however, current knowledge including genetic mapping of pigmentation variants for leaves, roots, and fruits (Albert *et al.*, 2015; Yan *et al.*, 2020; Xu *et al.*, 2020; Yu *et al.*, 2020) and ectopic expression of floral regulators in vegetative tissue (Albert *et al.*, 2020), indicates that the transcriptional MBW regulon is broadly involved in pigmentation patterning across diverse organs.

Our coleus breeding population expresses a tremendous diversity of pattern combinations. Rife and Boye (1938) recognized the potential of this prized ornamental, and proposed using Coleus as a model to dissect genetic regulators for color patterning. This suggestion did not get much traction, and we still know relatively little about color patterning in this unique ornamental. After 80 years of stalled progress, a renewed focus on the genetic regulation of pigmentation production and patterning would not only advance ornamental breeding, it would push the limits of Turing’s reaction diffusion model, reaching to describe the truly complex pattern variants that have drawn admiration from scientists and gardeners alike.

## Acknowledgements

We are grateful to the undergraduate researchers who assisted with data collection at the University of Florida, and to Joshua Tester for growing the plants. This work was funded by startup funds from the Donald Danforth Plant Science Center, and royalties from the UF Coleus Breeding Program. This project was also supported by the USDA National Institute of Food and Agriculture, and by Michigan State University AgBioResearch. MHF was supported by an NSF PGRP Postdoctoral Fellowship (IOS-1523668).

## Author Contributions

MHF, VC, and DHC designed the imaging pipeline, DC bred the Coleus population, MHF and VC collected the data, and ML developed and performed the HTP color analysis. All authors contributed to writing the manuscript.

## Supplemental Figures

**Figure S1:**
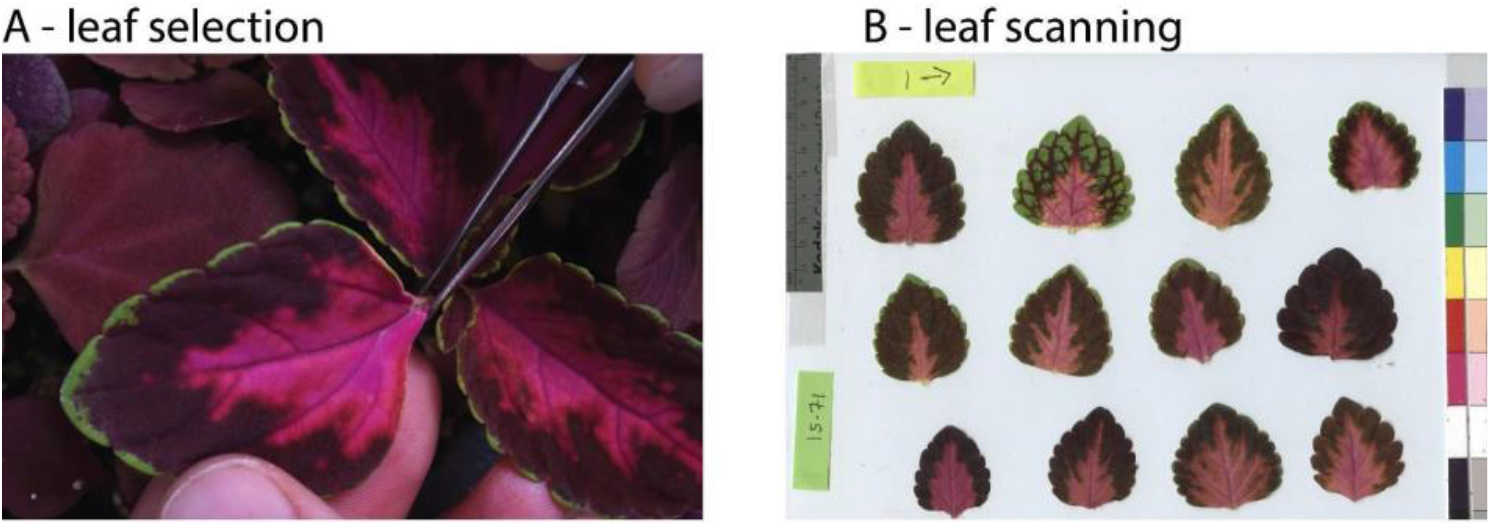
Data collection. **(A)** Data were collected for one leaf from the first fully expanded leaf pair. **(B)** Leaves were imaged on a flatbed scanner with a color card for color correction and a ruler.

**Figure S2:**
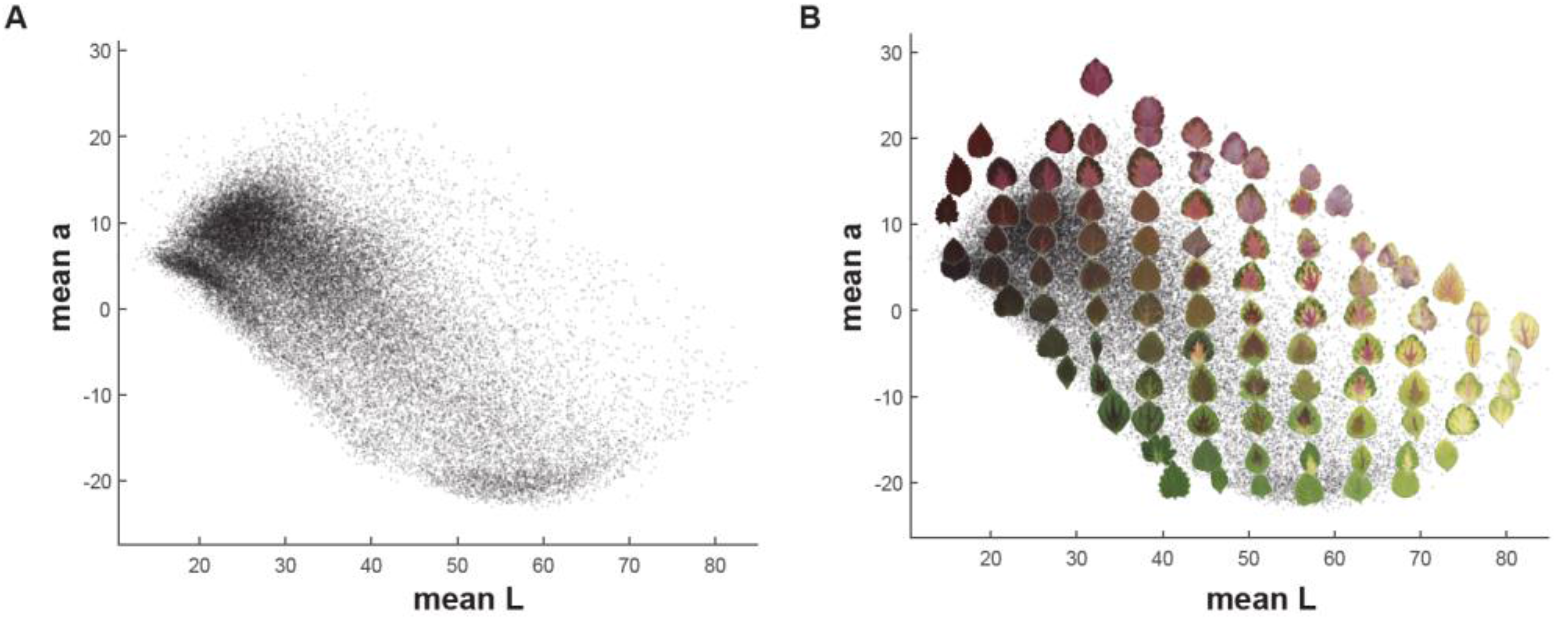
Color distribution for mean L and mean a. Plot of mean L (x-axis) and a (y-axis) for the full coleus population with point density (A) and example leaves overlaid on top of points (B).

**Figure S3:**
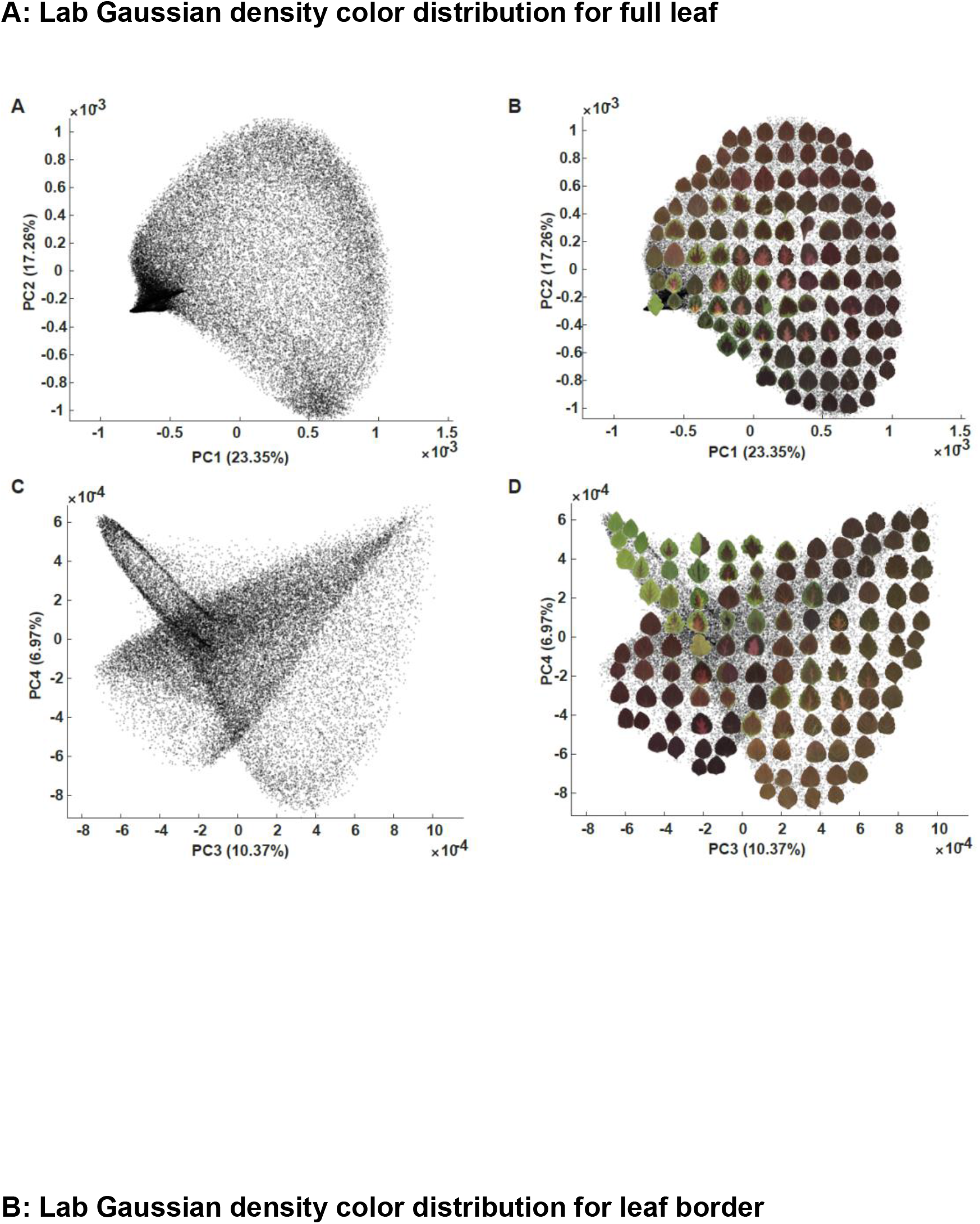
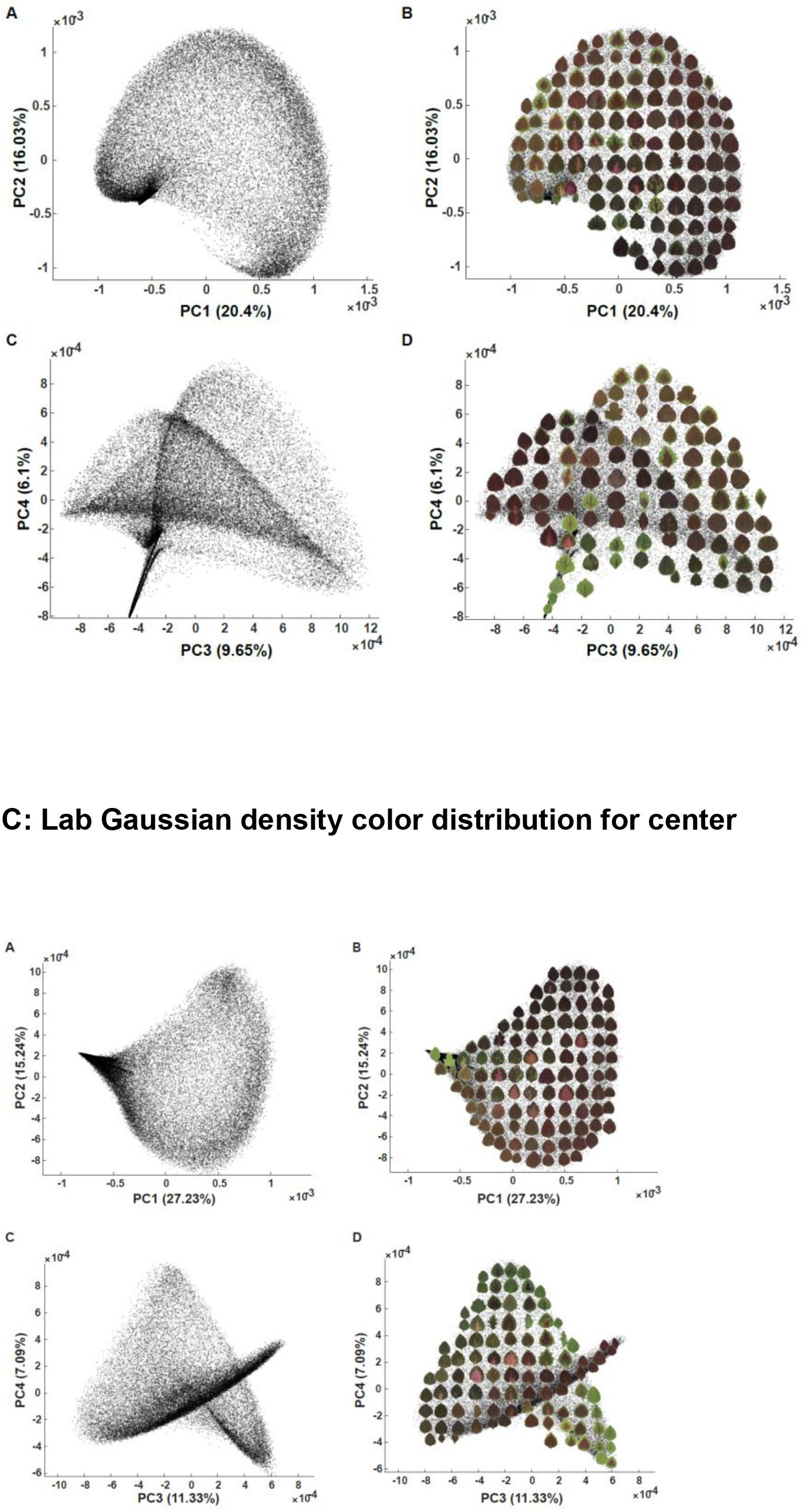
Principal Component Analysis for color in the entire coleus population using segmented or full leaf data sampling. PCA plots for Lab Gaussian density of color distribution with example leaves overlaid on top of data points for the full leaf (A), leaf border (B), and leaf center (C).

**Figure S4:**
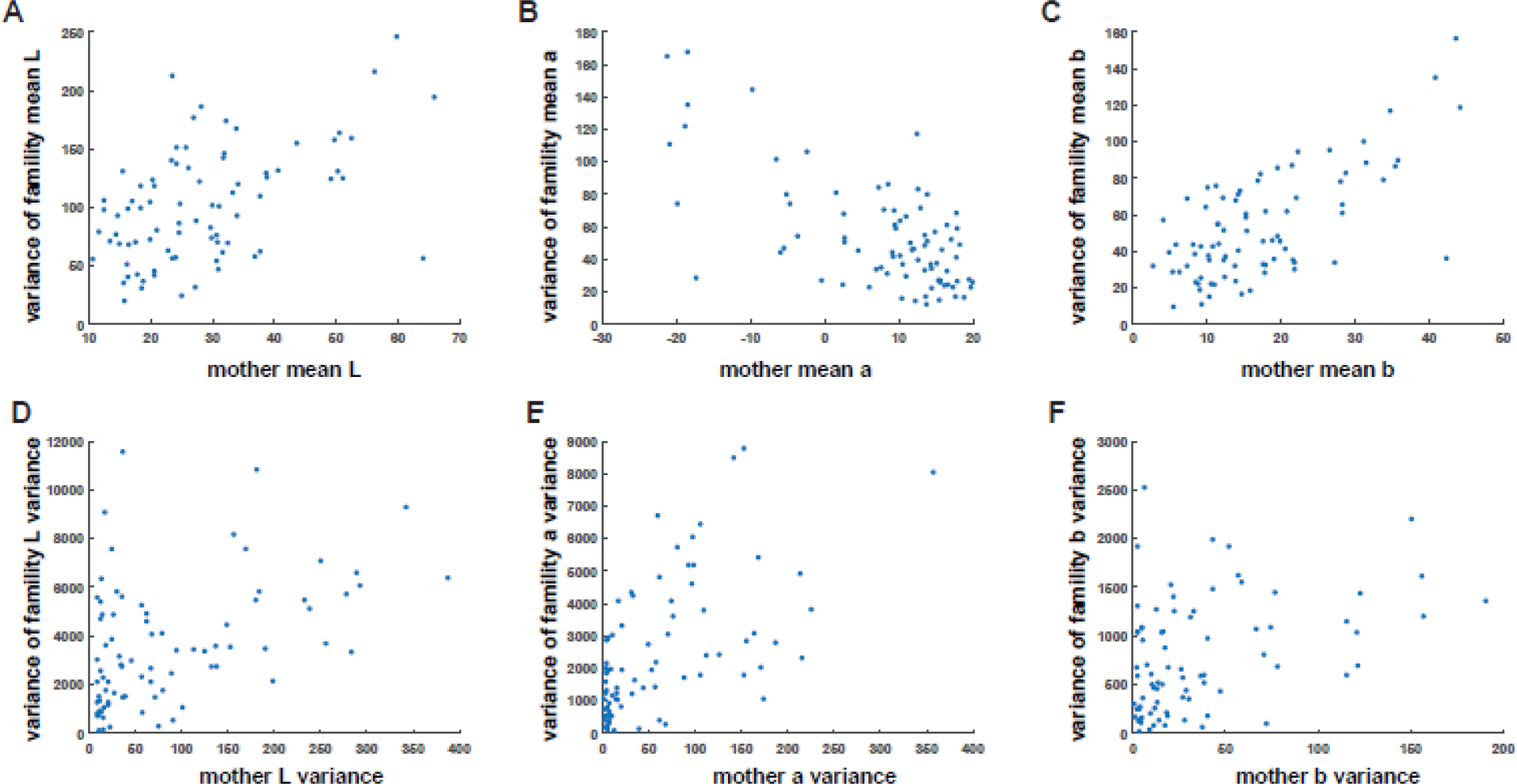
Maternal Plant-Progeny relationships based on mean and variance of Lab color. Scatter plots with x-axis represents the mean L (A), mean a (B), and mean b (C) of maternal plant and y-axis represents the variance of the corresponding mean value of the progeny, respectively; Scatter plots with x-axis represents the variance of L (D), variance of a (E), and variance of b (F) of maternal plant and y-axis represents the variance of the corresponding variance value of the progeny, respectively.

